# Synthetic organelles enable protein purification in a single operation

**DOI:** 10.1101/2024.05.17.594729

**Authors:** Haotian Guo, Jionglin Chen

## Abstract

Protein purification is crucial for various applications in life sciences research and biotechnological industry, yet traditional chromatography methods are often complex and suffer from limited scalability and sustainability. Here we report a biological approach utilizing synthetic organelles for protein purification, termed Program Architecture of Nano-Device Actions for Purification (PandaPure). PandaPure includes a biomimetic mechanism of sorting proteins of interest into organelles through signal sequence domains during expression, and subsequently releasing said proteins via cleavage of organelle-locating signals in a single operation. In the commonly-used expression host *Escherichia coli*, PandaPure enables high-yield, high-purity production of target proteins, and significantly reducing environmental impact. We demonstrate its scalability and versatility with different designs, scales, and across diverse proteins, promising wide-ranging benefits in research and industrial settings.

## Main Text

Protein purification serves as a fundamental process in both research and industrial applications within the life sciences, enabling advancements in fundamental biology and the development of biotechnological products (*1, 2*). However, conventional methods have long struggled with complexities related to scalability and sustainability (*3-6*). Chromatography-based techniques, relying on solid supports, dominate these methods, often leading to labor-intensive workflows with limited scalability (*3*). Tag cleavage, removal (*4*) and other downstream processing (*5*) require multiple rounds of chromatography, further posing the complexity. Moreover, concerns about chemical waste in purification processes highlight the need for more sustainable alternatives (*6*). While strategies like protein secretion (*7, 8*) and direct precipitation (*9*) offer some simplification, they often suffer from issues of low yield or purity. Consequently, there is a pressing demand for innovative approaches to protein purification.

To address these challenges, we introduce the Program Architecture of Nano-Device Actions for protein Purification (PandaPure), drawing inspiration from the efficient protein sorting, storage, and conditional release observed in secretory granules (*10*). By leveraging synthetic organelles (*11-15*), PandaPure offers a streamlined expression and purification procedure, where proteins of interest fused with a cleavable signal are spontaneously isolated by synthetic organelles, and can be recovered after cell lysis and a tag cleavage reaction (Fig. 1 and fig S1). Therefore, PandaPure condense the traditional workflow of affinity purification, tag cleavage and tag removal by a single operation of protein recovery.

**Fig. 1.**
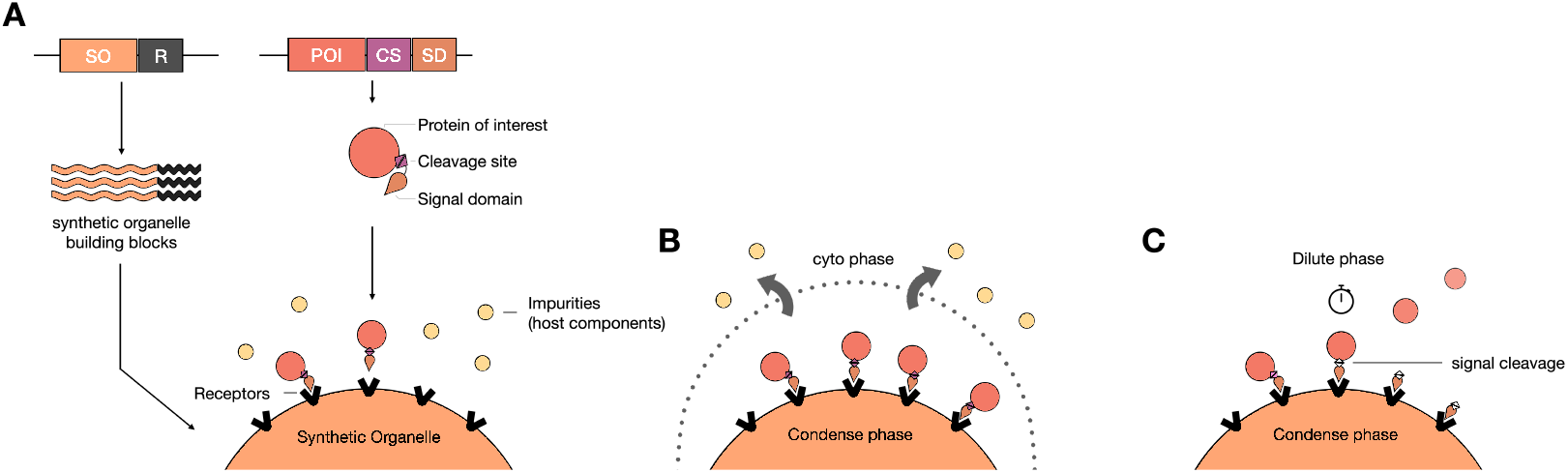
PandaPure design. (**A**) Coexpression of building blocks of synthetic organelles, and protein of interest fused to cleavable signal domain targeting the cognate receptor on organelles; **(B)** separation of dilute and condense phases to remove soluble impurity from host components; **(C)** protein recovery by self-cleavage of signal domain, to obtain tag-free proteins. SO. synthetic organelle building component, R, receptor, POI, proteins of interest, CS, cleavage site, SD, signal domain.

In this study, we provide a comprehensive exploration of the PandaPure framework, emphasizing its versatility, scalability, and sustainability. Through experimental demonstrations, we reveal PandaPure’s remarkable efficacy in achieving high yields and purity of desired proteins, using RNA synthetic organelles (*11*). Furthermore, we discuss the broader implications of PandaPure in diverse protein research and industrial applications, highlighting its potential to significantly enhance bioprocessing efficiency and sustainability.

### Validation of PandaPure Design Principle

For a proof of principle, we design a system according to PandaPure framework as the following. *Escherichia coli* is selected as the host organism, for its wide adaptation, high expression levels, and rapid growth in simple feedstocks, making it suitable for low-cost and sustainable protein production (*16*). For the synthetic organelle and protein sorting system, we use synthetic RNA organelle described in a previous work termed transcriptionally engineered addressable RNA solvents (TEARS) (*11*), where tandem repeats of triplet nucleotides undergo liquid-liquid phase separation (LLPS) to form synthetic organelles, and widely-adapted MS2 tracking system (*17*) serves as the recruiting receptors. Cognate signal domain is tandem dimeric MS2 coat protein (MCP) fused with the protein of interest. It is expected that protein expression, recruitment and enrichment occur simultaneously during the growth and expression. Then, following cell disruption, soluble impurities (dilute phase) can be easily removed from cell debris containing the organelles (condense phase) by centrifugation or other separation techniques. For the visualization of PandaPure workflow, we choose superfolder green fluorescent protein (GFP) (*18*) as the protein to be purified. Between GFP and MCP, we place a cleavage site (Fig. 1A). We hypothesize that once the signal domain is cleaved, organelles will spontaneously exclude the target proteins into dilute phase. Then, the target proteins (dilute phase) and organelles with remaining signal domains (condense phase) can be easily separated by brute forces, enabling the recovery of highly pure proteins in a single centrifugation. Such site can be the substrate of endoproteases (*4*) or an intein domain (*19, 20*). Here we apply a system that cut serine residue by catalysis of nickel, given the benefits of simplicity for demonstration (*21-23*).

To assess the performance of synthetic organelles in PandaPure, we express the systems in BL21(DE3) strains under the induction of 0.5 mM isopropyl β-D-1-thiogalactopyranoside (IPTG) and 150 ng/ml anhydrotetracycline (aTc) in 50 ml culture at log phase for 16 hours (see details in Methods). Overnight cells are harvested by centrifugation. Remarkably, a clear-cut demixing of differentiated cells occurs (Fig. 2A), implying that TEARS-GFP continuously grew in the mother cells and made them denser than daughter cells (Fig. 2B), reckoning that transmission electron microscopy showed a higher electron density of TEARS and the fluorescent imaging of bacteria in mother machines (*11*).

**Fig. 2.**
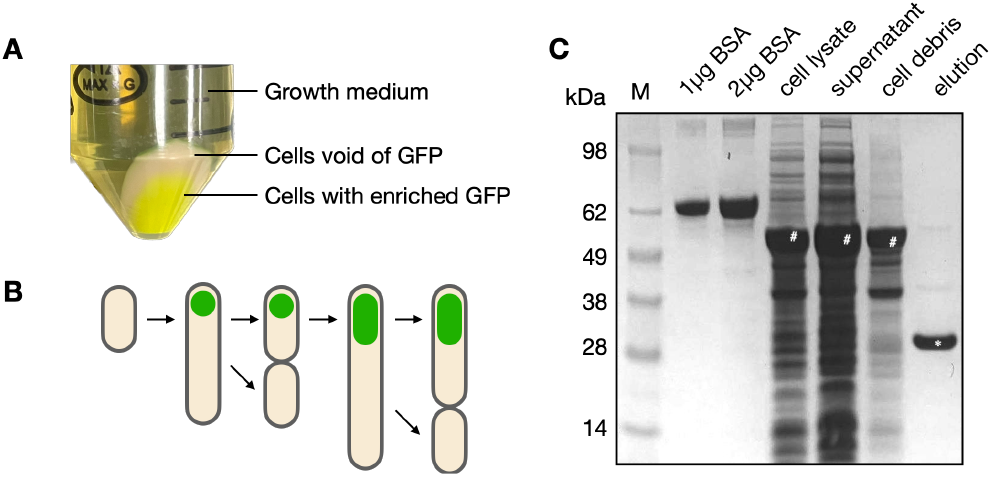
PandaPure of GFP. (**A**) Recruitment of GFP into organelles drives differentiation of bacterial cells, causing density differences, leading to separation after centrifugation. (**B**) the diagram of differentiation of bacterial cells. Mother cells accumulates protein of interests in growing synthetic organelles (in green) and becomes denser than daughter cells. (**C**) Organelles remain intact in the cell debris after the detergent-based cell lysis treatment. (**D**) PandaPure enables recovery of highly purified GFP. M, protein weight marker; BSA, bovine serum albumin; cell lysate, whole-cell sample after detergent and lysozyme treatment; supernatant, dilute-phase supernatant of cell lysates; wash, supernatant of the washing; cell debris, condense-phase pellet of cell lysates; elution, recovered dilute-phase supernatant after signal cleavage reaction.

Then, we lyse bacterial cell to separate soluble impurity in the dilute phase, i.e. supernatant, and synthetic organelles in the condense phase, i.e. cell debris. It was suggested that liquid-phase structures dissolve under rapid dilution (*11, 24-26*). However, here we report that under the treatment with non-ionic detergent, lysozymes and even endonuclease, TEARS surprisingly maintain its function to sequester the MCP-tagged GFP in the cell debris (Fig. 2C). It is likely that surfactants enhance the separation of those compartments, consistent with previous studies of LLPS during amphiphilic self-assembly (*27*).

After, we wash the cell debris for once, and resuspend it for the reaction of signal domain cleavage overnight. Centrifugation of the reaction mixture separates target protein in dilute phase, i.e. elution, and remained organelle components in the condense phase. Visualized by denatured electrophoresis, we demonstrate that GFP is purified without undesired proteins, or only occasional leaky release of MCP-GFP proteins (Fig. 2C, and fig. S2).

### Simplified PandaPure Enables Minimal Operations in a Single Reaction Vessel

Next, we aim to further explore the limits of simplicity by removing washing, and the addition of lysozyme during lysis which causes a significant increase of the viscosity and impairs the easiness of following operations (Fig. 3A). Our previous workflow only applies one-time washing and results in high purity. Surprisingly, we show impurities are only increased at a marginal level without washing (Fig. 3B). Without lysozyme treatments, similar amounts of GFP are purified, but at a lower purity as expected (Fig. 3B). Therefore, the cell lysis efficiency is a major determinant of the purity of PandaPure products. It is implied that TEARS isolate target proteins of interest in vivo at high efficiency so that PandaPure can tolerate suboptimal cell lysis.

**Fig. 3.**
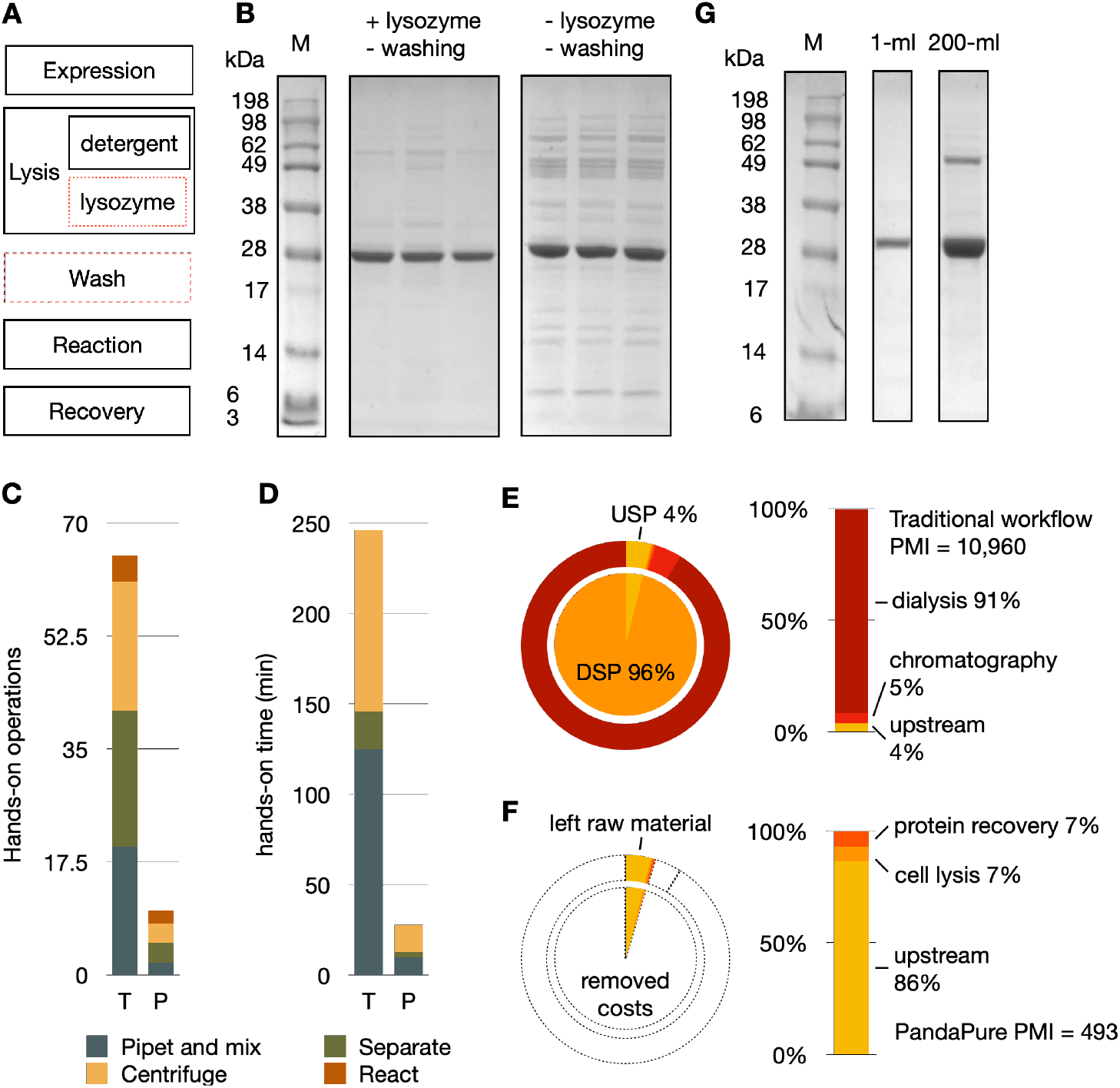
Simplicity, sustainability and scalability of PandaPure workflow. (**A**) Simplifying PandaPure workflow by removing lysozyme treatment and washing (in red dash-line box). (**B**) Simplified PandaPure workflows results in high yield, and lower but comparable purity. (**C and D**) PandaPure reduces the complexity of operations counted in basic units (**C**), and hands-on times (**D**), counting from the harvest of expressed cells. T, traditional workflow, including cell harvest, lysis, affinity step, tag cleavage and tag removal. P, simplified PandaPure workflow, including cell harvest, lysis, and protein recovery (fig. S3). (**E and F**) PMI analysis (kilogram of waste per kilogram of protein product) for the production of GFP by using traditional workflow combining chromatography, dialysis and protease treatment (**E**), versus the simplified PandaPure workflow (**F**). The pie charts show how each stage of bioprocessing contributed to the overall PMI, normalized by traditional workflow. The bar graphs show a breakdown of mainly-used materials. (**G**) PandaPure scales from 1 ml in microplates to 200 ml bacterial culture in flasks.

Remarkably, the simplified PandaPure without either washing and lysozymes still results in a comparable purity to many batch-bind affinity purifications. We anticipate this simplified version of PandaPure can be efficient enough for a variety of applications.

Thus, PandaPure enables the least effort to obtain purified tag-less protein from cell cultures up to date. Considering breaking down the workflow into basic units, (a) mixing, (b) centrifugation, (c) separating supernatant and pellets, and (d) incubating reactions, the simplified PandaPure only contains 10 units starting from cell harvest. In comparison, a traditional workflow that involves affinity purification and protease treatment has 65 units (Fig. 3C, and fig. S3). By estimation, PandaPure only require hands-on operations for 30 minutes, 8-fold less than the traditional workflow (Fig. 3D). Moreover, PandaPure is a “one-pot” purification procedure in the sense that all operations can be carried out in a single reaction vessel.

### Enhancing Environmental Sustainability in Protein Purification

Using the metrics acquired from our 50-ml culture, we performed a process mass intensity (PMI) analysis to evaluate the material efficiency of PandaPure (Fig. 3E and 3F). Even without additional optimization for the costs and wastes, PandaPure has already surpassed a traditional workflow combining batch-bind affinity chromatography, protease and dialysis by 20-fold. Per kg GFP, PandaPure requires 493 kg of raw materials, compared 11,000 kg from a traditional workflow. This improvement is primarily attributed to the extreme simplification of downstream processing, substantially reducing the economic and environmental costs associated with protein production. 96% of raw materials are cost in the dialysis and chromatography in a traditional workflow; in contrast, only 14% of materials are used to lyse cells and recover proteins in a simplified PandaPure workflow. Further enhancements are feasible by optimizing protein expression levels to minimize the overall volume of bioprocessing in future endeavors of improvements in sustainability metrics.

### Scalability of PandaPure

Scalability, either scaling up or down in volumes, or scaling in or out in throughput, is a critical aspect in protein production. Previously, a variety of materials and methods were developed for chromatography to adapt different applications, such as magnetic beads for small volumes used in high-throughput screening and automation, and radical flow columns for large volume production. Instead, PandaPure provides a universal solution from different volumes and throughputs as the purification occurs in vivo. Despite the inherent differences in expression titers and leaky releases, PandaPure can apply for 1 ml culture in parallel in 96-well deep-well microplates used for high-throughput screening to 200 ml culture in 1L flask for small-scale production (Fig. 3G). Thus, we anticipate there is a potential for PandaPure to apply for diverse applications from early-stage discovery to industrial production in the future.

### Improving the production of industrial and therapeutical proteins with PandaPure

Finally, we demonstrate PandaPure’s capacity to address diverse bioproduction challenges, by focusing on a selected group of proteins that are renowned for their commercial values and also their production challenges (table S1).

First, we validate our approach with tool enzymes, including T4 DNA ligase for molecular cloning (*28*) and a variety of applications (*29*), type II-S enzyme BsaI for scarless cloning (*30*) and mRNA vaccine production (*31*), and methyl-directed endonuclease DpnI often applied to remove host templates, such as in site-directed mutagenesis (*32, 33*) (Fig. 4A to 4C). Notably, host DNAs and related physiologies are susceptible to these enzymes, thereby asking for specific strategies to produce them, including fed-batch fermentation (T4 DNA ligase, *34*), applying restriction-modification (R-M) system (BsaI, *35*), and dam/dcm deficiencies (DpnI, *36*). As synthetic organelles can decouple host physiology and recombinant protein expression by sequestering the proteins of interest (*37*), PandaPure offers a streamlined expression and purification route, eliminating the research and development for specific modifications, and presenting a unified solution to both expression and purification hurdles.

**Fig. 4.**
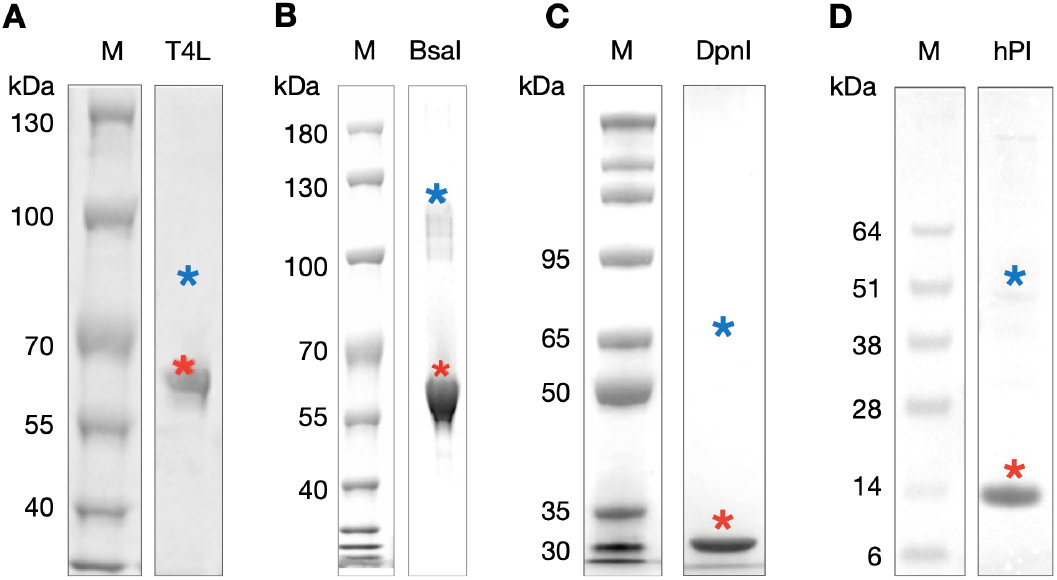
SDS-PAGE of PandaPure products with commercial values. Proteins of interest (POI) includes (**A**) T4 DNA ligase (T4L), (**B**) BsaI, (**C**) DpnI, (**D**) human proinsulin (hPI). M, protein markers. Asteroids, expected molecular weights of tag-less POI (red), or the leaky release of tagged POI (blue).

Then, we showcase PandaPure’s potential for therapeutics, with human proinsulin (hPI), the precursor vital for producing recombinant insulin (*38, 39*). Typical expression of hPI produces inclusion bodies (*38*) in *E. coli* because disulfide bond cannot form in its reductive cytoplasm (*40, 41*). Traditionally, refolding proteins and multi-step purification are necessary in downstream processing (*42*). Leveraging improved disulfide bond formation and protein folding in TEARS (*43*), we condensed the production of hPI into a single efficient step (Fig. 4D). This innovation not only accelerates production by days, but also drastically reduces costs and environmental impacts by eliminating refolding.

## Conclusion

In conclusion, PandaPure offers a promising approach to protein purification, leveraging synthetic organelles to sort, purify and store target proteins specifically, and the self-cleavage reaction of signal domains to release target proteins efficiently. This paradigm shift, from traditional chromatography and secretion-based methods to a biological approach, brings scalability, sustainability, and simplicity to the forefront.

PandaPure represents currently the simplest workflow to obtain purified, tag-less proteins. It largely reduces the manual work in protein purification. Replacement of reaction-centrifugation methods by other reaction-separation techniques may further reduce the complexity. We anticipate it can promote a wider adaptation of recombinant protein production in the fundamental research. Moreover, the elimination of solid supports used in chromatography makes PandaPure a generic method, spanning across small-volume production such as high-throughput screening to large-volume bulk production, further lowering the learning curves.

For production, demonstrations of GFP, tool enzymes, and proinsulin underscore PandaPure’s potential for low-cost, sustainable, and scalable protein production. The economic and environmental costs are considerably reduced compared to the traditional workflows, especially for ones that are difficult to express. Unlike chromatography, PandaPure’s synthetic organelles boast immense capacity (*11*), extending beyond protein expression yield limitations.

While tested primarily in *E. coli*, PandaPure holds promise for application across a wide range of organisms. Overall, PandaPure represents a transformative tool with far-reaching implications for the future of protein production and manufacturing, offering a streamlined, sustainable, and versatile approach to meet the evolving demands of bioprocessing.

## Acknowledgments

We thank members of INSERM U1001 (now U1284, France) for their pioneering work on protein purification using synthetic organelles, by H. Guo, J. Ryan, V. Pabst, and A. B. Lindner. While their efforts only achieved biocondensate purification but not protein purification (*11*), it provided valuable insights that we have incorporated into this work. We are grateful to the Ailurus team for helps in developing PandaPure method, with thanks to E. Qiu, J. Shan for early validation efforts, N. Jiang, J. Yang, X. Wang for further repeating PandaPure of GFP, where N. Jiang’s repeats are used in this manuscript (Fig. 2D, and fig. S2B). We thank J. Hu, A. Hao for reviewing the work and provide comments.

## Author contributions

Conceptualization: H.G.

Methodology: H.G.

Investigation: H.G. (Fig. 1-3), J.C. (Fig. 4)

Project Administration: H.G.,

Funding Acquisition: H.G.

Writing - Original Draft: H.G.

## Data and materials availability

All materials are commercially available from Ailurus at https://www.ailurus.bio/. All data are available in the main text or the supplementary materials.

## Supplementary Materials

Materials and Methods

Figs. S1 to S3

Tables S1

