## Supplementary material for "Synthetic organelles enable protein purification in a single operation": Materials and Methods, Figs. S1 to S3, Tables S1

**The PDF file includes:**

Materials and Methods
Figs. S1 to S3
Tables S1

### Materials and Methods

Strain, growth medium and conditions.

The *E. coli* strain BL21(DE3) (NCBI Taxonomy ID: 469008) and BL21AI (Invitrogen) were used for PandaPure validation, protein expression and purification. Plasmids are transformed into chemical competent cells by heat shock. For transformation of more than two plasmids, no more than two plasmids were transformed first to create an intermediate strain, and then the rest of the plasmids were transformed into the intermediate strain, no more than two plasmids per generation. Unless specified otherwise, 150 ng/mL anhydrotetracycline (aTc), 0.5 mM isopropyl β- d-1-thiogalactopyranoside (IPTG) were used for induction; 100 mg/mL ampicillin, 50 mg/mL kanamycin, 100 mg/mL spectinomycin and 30 mg/mL chloramphenicol were used for antibiotic selections; Super Optimal broth with Catabolite repression (SOC) medium were used for growing bacteria in seed cultures, Luria-Bertani (LB) broth Miller recipe were used for protein expression; bacteria cultures were incubated at 37°C and shaken at 220 revolutions per minute (rpm) for tubes, flasks, and 900 rpm for microplates.

Plasmid and cloning methods

PandaPure Protein Constructs include (1) one plasmids encoding synthetic organelles and (2) second plasmid encoding expression of proteins of interest fused with cleavable signal domain, respectively, and are used to validate PandaPure workflow (table S1). Plasmids newly constructed in this study are assembled by DNA manufacturers, through inserting de novo synthesized gene fragments into designated PandaPure expression vector (PPC) with seamless cloning methods.

Protein expression

Bacteria were first prepared through a process to homogenize the bacterial physiology. For each experiment, a new transformation is performed. Unless specified otherwise, we do not start experiment from glycerol stocks, to avoid any potential bacterial degeneration. Single colonies were inoculated into a seed culture, and incubate overnight. Then overnight cultures were diluted 200 or 250-fold in LB medium (initial OD600 is about 0.05), and grown back to the mid-exponential phase in about 2 hours. Then IPTG and aTc were directly supplied to induce the expression of components that form synthetic organelles and the expression of proteins of interest, for extra hours.

Cell lysis and isolation of synthetic organelles

Induced cultures were centrifuged at max speed for 5-10 minutes. Supernatant were discarded and the pellets were resuspended in PandaPure Lysis Reagent at 8 ml per gram biomass, or 1/5 of the culture volume for consistencies in microplates, or other non-ionic detergents such as B-PER™ Bacterial Protein Extraction Reagent (Thermo Fisher Scientific), BugBuster® Protein Extraction Reagents (Merck Millipore), under the instructions of manufacturers. Lysozyme (Sigma Aldrich, or Beyotime), and Benzonase® Nuclease (Sigma Aldrich) are used for more through lysis. Cell lysis is performed by incubating the reaction at room temperature or 37°C for higher efficiency in rotators or shakers, for 15-20 minutes. Cell lysates are centrifuged at max speed for 10 minutes. Supernatants (dilute phase) are discarded and the pellets (condense phase) containing synthetic organelles are kept for further steps or analysis.

Protein recovery from synthetic organelles

Condense-phase pellets are resuspended in PandaPure Protein Reagent at the same volume of previously used lysis reagent, and incubate at 37°C for higher efficiency in rotators or shakers, overnight to 24 hours. Reaction mixtures are centrifuged at max speed for 10 minutes. Supernatants (dilute phase) are kept for further steps or analysis, and the pellets (condense phase) are discarded.

Directed evolution of plasmids encoding TEARS system

To isolate more stable plasmid systems for TEARS, we grow bacteria strain DH5alpha transformed with the original TEARS design previously described (*11*) on LB agar plates with chloramphenicol antibiotic at 4°C for months. Then we streak the old colonies into a fresh LB agar plate, to isolate single colony that maintains the plasmids that are evolutionary stable. Plasmids are then extracted for Sanger sequencing. Candidates that still encode TEARS are kept. Plasmids are retransformed into DH5alpha. Then we repeat this process, until the TEARS plasmid remains stable.

SDS-PAGE gel electrophoresis

20 μL of samples are denatured by adding 5 μL loading buffer at 95°C for 10 min. 10 μL denatured samples are loaded in precast SDS-PAGE gels. After electrophoresis, gels are stained to visualize protein bands, manually, or using eStain L1 Protein Staining Device (Genscript).

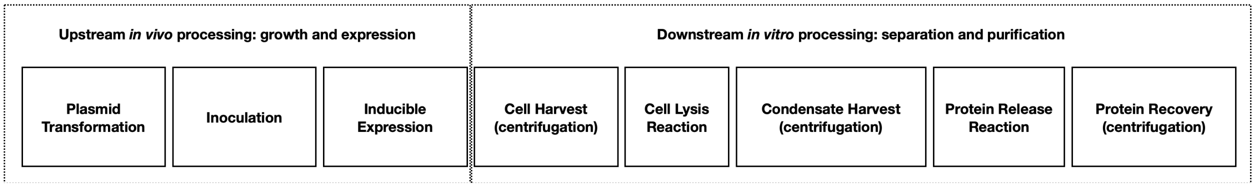


**Fig. S1. The workflow of PandaPure.**

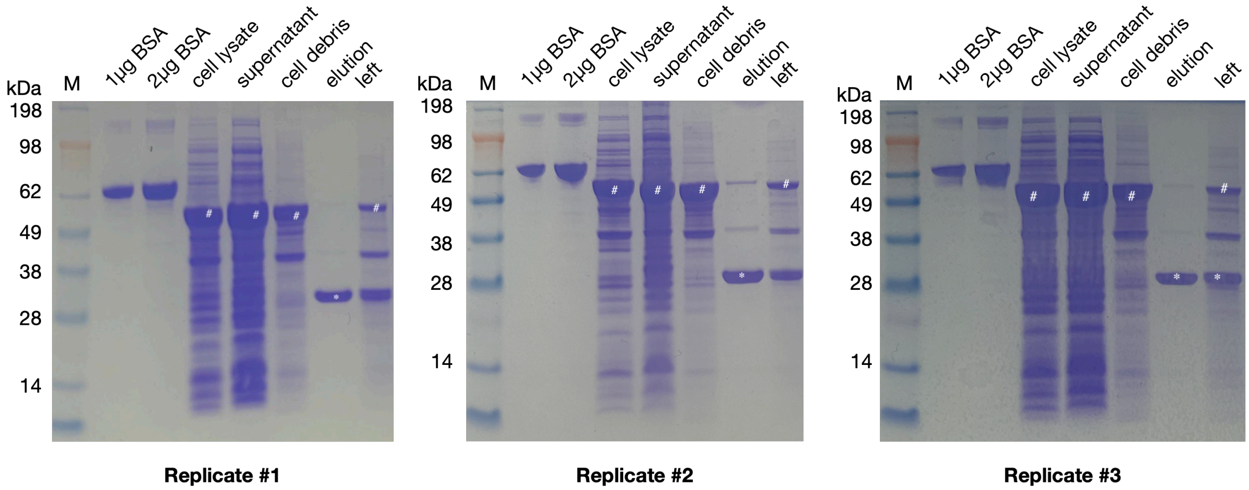


**Fig. S2. Raw images of SDS-PAGE gel electrophoresis: PandaPure of GFP with TEARS.** (SDS-PAGE of PandaPure workflow in three biological replicates (different bacteria colonies). M, protein weight marker; BSA, bovine serum albumin; cell lysate, whole-cell sample after detergent and lysozyme treatment; supernatant, dilute-phase supernatant of cell lysates; wash, supernatant of the washing; cell debris, condense-phase pellet of cell lysates; elution, recovered dilute-phase supernatant after signal cleavage reaction.

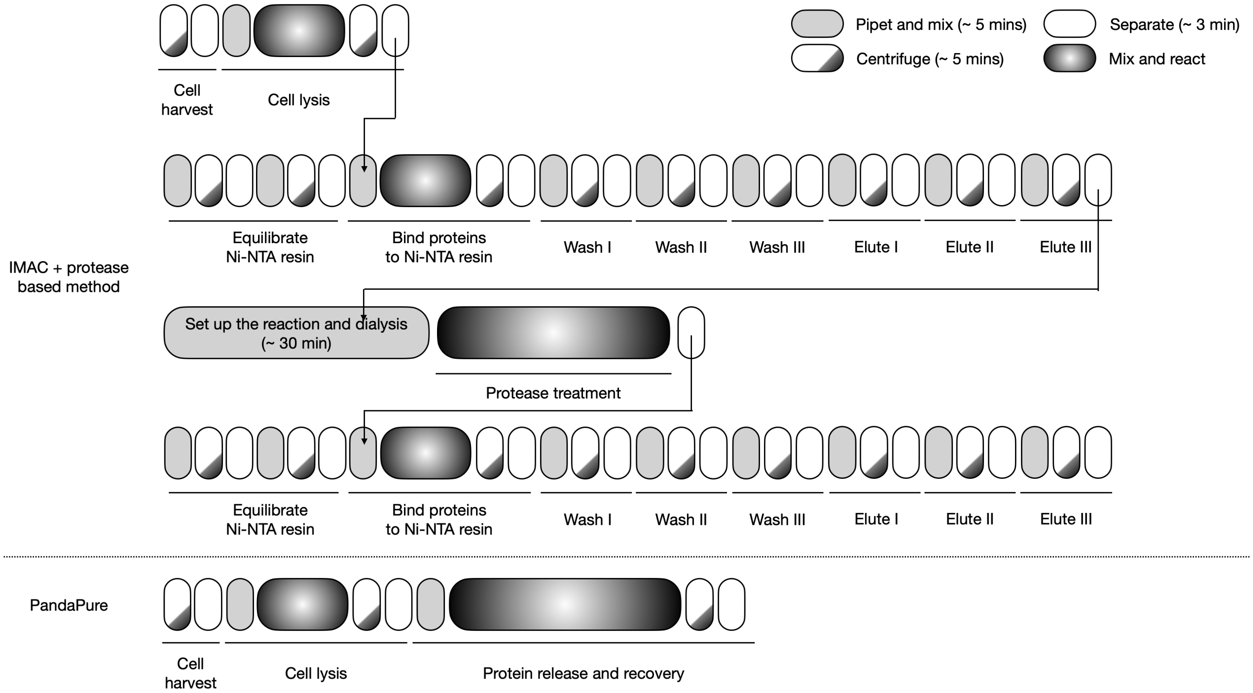


**Fig. S3. Workflow comparison between traditional workflow using immobilized metal affinity chromatography (IMAC) and protease treatment, versus PandaPure.** Representative data about the operations for traditional workflow are acquired from HisPur™ Ni-NTA Resin (Thermo Fisher Scientific).

**Table S1. Plasmids used in this study**

| Name | Source | Description | Related Figure |
| --- | --- | --- | --- |
| pTEARS | (*11*) | 47 CAG - 12 MS2, expressed under PLtetO-1 promoter | Fig. 2C |
| pTEAR2 | This study | 72 CAG - 12 MS2, expressed under PLtetO-1 promoter | Fig. 2A, and 2D, fig. S2B, Fig. 3B, and 3G |
| pIG-K133 | (*17*) | Tandem dimer MS2 coat protein (tdMCP) – GFP mut3 fusion | Fig. 2C, and fig. S2A |
| pPPN | This study | PandaPure N-term expression vector, with cloning site for POI, fusing tdMCP and cleavage site at N-terminal of POI, expressed under T7-lac promoter | Intermediate plasmid |
| pPPC | This study | PandaPure C-term expression vector, with cloning site for POI, fusing cleavage site and tdMCP at C-terminal of POI, expressed under T7-lac promoter | Intermediate plasmid |
| pPPC-sfGFP | This study | Superfolder GFP cloned in PandaPure expression C-term vector | Fig. 2A, and 2D, fig. S2B, Fig. 3B, and 3G |
| pPPC-T4 DNA ligase | This study | Hi-T4 DNA ligase (New England Biolab) cloned in PandaPure expression C-term vector | Fig. 4A |
| pPPC-BsaI | This study | BsaI cloned in PandaPure expression C-term vector | Fig. 4B |
| pPPC-DpnI | This study | DpnI cloned in PandaPure expression C-term vector | Fig. 4C |
| pPPN-hPI | This study | Human proinsulin in PandaPure expression N-term vector | Fig. 4D |
